# Circular RNA - circCLIP2 is predominantly expressed in GSC niche and enhances glioblastoma aggressiveness via EMT and ECM signaling

**DOI:** 10.1101/2024.12.01.626287

**Authors:** Julia Misiorek, Żaneta Zarębska, Konrad Kuczyński, Julia Latowska-Łysiak, Adriana Grabowska, Paweł Głodowicz, Marcin Piotr Sajek, Anna Karlik, Dorota Wronka, Anna Maria Barciszewska, Łukasz Przybył, Katarzyna Rolle

## Abstract

Glioblastoma (GBM) is the most common and malignant type of brain tumor in adults and no effective therapies exist to combat this disease. Surgical resection, radio- and temozolomide-based-chemotherapy are insufficient mainly due to inter- and intra-tumoral heterogeneity of GBM tumors, therefore personalized approaches are required. Additionally, diverse tumor microenvironment with a leading role of glioblastoma stem cells (GSCs) in hypoxic core underlie the aggressiveness and recurrence of GBM. However, the molecules and pathways which directly promote tumor aggressiveness together with the driving process of epithelial-to-mesenchymal transition (EMT) and the components of extracellular matrix (ECM), remain only partially covered with prior studies focused mainly on coding oncogenes and tumor suppressor genes. Over the past years, a special attention has been paid to non-coding, circular RNAs (circRNAs) differentially expressed in various cancers, including GBM. Despite high abundance and stability, the exact biological roles of most individual circRNAs have not been fully revealed, so far. Besides anti-cancer targets, circRNAs have been considered as essential biomarkers and stratifying criterion of malignancy. Thus, studies on functions of differentially expressed circRNAs in GBM are an emerging field to be explored. In herein study, the biological role of circRNA CLIP2 (circCLIP2) was found highly overexpressed in primary tumor tissues. Upon the knock-down of circCLIP2 in 2D and 3D *in vitro* models, functional tests performed clearly indicate a significant contribution of circCLIP2 to aggressive potential of GBM as showed by decreased rates of proliferation, migration and invasion. Additional studies indicated predominant expression of circCLIP2 in GSC fractions additionally elevated by hypoxic conditions. The increased rates of proliferation and migration as well as molecular analyses were also confirmed in *in vivo* settings where xenograft model was applied. The expression of EMT marker *Snail2* and ECM-degrading enzyme metalloproteinase 9 (*MMP9*) were significantly downregulated in the tumors formed from circCLIP2 silenced cells. Taken together, these results indicate a strong contribution of circCLIP2 in the aggressive phenotype of GBM which is due to a modulation of GSC proliferative potential as well as migration and invasion via EMT together with the modulator of ECM composition.

## 1. Introduction

Cellular potential of unrestricted proliferation and invasion are key determinants of tumor aggressiveness. To the most aggressive types of primary tumors belongs glioblastoma (GBM) which additionally displays a high intra- and inter-tumoral heterogeneity, the factor that determines resistance to chemio- and radiotherapy against GBM. [1] There are no molecular biomarkers of diagnostic and prognostic relevance that could consider GBM aggressiveness as determinant factor. The only verified genetic biomarker of GBM malignancy which distinguishes glioblastoma from astrocytoma is the presence of mutation in genes encoding the cytoplasmic and mitochondrial enzyme isocitrate dehydrogenase (IDH1 and IDH2, respectively). [2] GBM tumors are highly recurrent which is fueled by the presence of glioma stem cells (GSCs) with self-renewal ability and multi-directional differentiation potential. Interestingly, it has been suggested that stem cells can acquire mesenchymal phenotype in the course of epithelial-to-mesenchymal transition (EMT) which facilitates cancer spread while sustaining pluripotency. [3, 4] From the molecular perspective, the discovery of crucial molecules for GBM development and progression is still the subject of extensive studies which focus on both coding and non-coding transcripts. CircRNAs are non-coding and highly expressed molecules in human brain. They are characterized by a covalently closed loop generated by non-canonical back splicing. Due to the absence of 5’ and 3’ ends, circRNAs are highly resistant to degradation by nucleases, which makes them exceptionally stable molecules. High stability and altered expression under pathological conditions render circRNAs as suitable biomarkers of diseases. Thus, selected circRNAs of well-studied biological relevance could serve as novel markers of diagnostic, prognostic or even therapeutic potential. [5] So far, the involvement of some specific circRNAs has been already reported in cellular processes including proliferation, apoptosis, migration, invasion and angiogenesis which manifest during GBM [6–9] circCLIP2 is derived from linear *CLIP2* gene which displays a brain-specific expression and encodes cytoplasmic linker proteins facilitating the binding of membranous organelles into microtubules. So far, the increased survival rate of patients suffering from GBM was found to correlate with low expression of *CLIP* genes. [10] Interestingly, the fusion of *CLIP2* with *MET* gene was reported in other type of brain cancer – glioneuronal tumor. [11] It has been observed that gene fusion triggers mitogen-activated protein kinase (MAPK)-signaling pathway which in turn affects cell migration, proliferation and progression of GBM. [11] Although, the function of linear *CLIP2* has been already established, the role and relevance of its circular form – circCLIP2 in GBM remains elusive.

To determine the possible function of circCLIP2 for the biology of GBM, we established the expression profile of CLIP2 in the GBM tissues confirming its overexpression. The functional analyzes upon siRNA-mediated knockdown show circCLIP2 contribution to the aggressive potential of GBM exhibited by decreased rates of proliferation, migration and invasion. Further, our studies also correlate circCLIP2 function with GSCs population and hypoxic niche. Finally, we confirmed circCLIP2 impact on the GBM progression in animal study. GBM xenograft-based analysis indicate that cirCLIP2 is involved in the EMT process and leads to the gene expression deregulation with the predominant knockdown of ECM-degrading enzyme metalloproteinase 9 (*MMP9*).

## 2. Materials and methods

### 2.1 Patient sample collection

The tumor tissues were received from the Department and Clinic of Neurosurgery and Neurotraumatology at the University of Medical Sciences in Poznan and the Department of Neurosurgery at the Multidisciplinary City Hospital in Poznan, both localized in Poland. Prior the experiments, the approval from The Bioethics Council of University of Medical Sciences (approval # 46/13) and patients’ consents were obtained. First, the tissues of primary GBM tumors (n=12) from patients were collected into MACS Tissue Storage Solution (Miltenyi Biotec) by surgical excision and next, transported within 1 h into research laboratory were RNA was isolated. The *“First Choice Human Brain”*, commercially available pool (n=23) of total RNA isolated from healthy human brains was purchased (Ambion, cat. No# AM6050) and utilized as control. Characterization of the tissues used for RNA isolation and the basic anonymized information from GBM patients is presented in Supplementary Table 1.

### 2.2 Cell culture

Human glioblastoma cell lines U251-MG and U138-MG were purchased from American Type Culture Collection (ATCC) while U-87 MG from Merck (Sigma). Cells were grown in Eagle’s Minimal Essential Medium (EMEM, Corning) or Dulbecco’s modified Eagle’s medium (DMEM, ATCC) as recommended by the manufacturers, supplemented with 1% penicillin– streptomycin antibiotic (Sigma-Aldrich) and 10% fetal bovine serum (FBS, Sigma-Aldrich). All cell-lines had adherent features while the U-87 MG cell line was partially (30%) forming spheres spontaneously under our standard culture conditions. Cells were grown at 37°C, in 95% humidity, and 5% CO_2_ conditions.

### 2.3 Spheroids formation

Human glioblastoma cell lines U138-MG and U251-MG served as material for generation of 3D structures - spheroids. To form the spheroids, 3 × 10^3^ cells/mL were seeded into a non-adherent 96 U-bottom plate in 200 µL of medium recommended by the manufacturer, supplemented with FGF and EFG growth factors, 1% penicillin–streptomycin antibiotic (Sigma-Aldrich). Cells were centrifuged at 300 x g for 3 min and grown into spheroids for 4 days in the incubator under standard culture conditions.

### 2.4 RNA extraction and quality validation

Total RNA was extracted from GBM patient tissue and GBM cell lines using the TRIzol reagent (Invitrogen) according to the manufacturer’s instructions. The DNase I treatment was applied on RNA samples using ready-to-use DNA-free^TM^ DNA Removal Kit reagents according to the manufacturer’s protocol (Ambion). The quality of purified RNA was determined by NanoDrop 2000 spectrophotometer (Thermo Fisher Scientific) and electrophoretic separation in 1.5% agarose gel.

### 2.5 Reverse transcription and qRT-PCR

The reverse transcription reaction was performed using the Transcriptor High Fidelity cDNA Synthesis Kit (Roche) according to the manufacturer’s protocol utilizing 500 ng of RNA. The real-time PCR reaction was performed in three technical replicates using the CFX Connect Real-Time PCR Detection System (Bio-Rad), LightCycler® 480 SYBR Green I Master (Roche), and primers designed using Primer-BLAST tool [27] (Thermo Fisher Scientific). The sequences of primers used in qRT-PCR analysis presents Supplementary Table 2. Hypoxanthine phosphoribosyltransferase (*HPRT*) gene was used as an endogenous control.

### 2.6 Sanger sequencing

CircCLIP2 was amplified by applying the following PCR protocol: 3 min of denaturation and repeated in 38 cycles: 15 sec of denaturation at 95°C, 30 sec of annealing at 63°C, and 20 sec of extension. The PCR products were visualized in 1% agarose gel for the quality check and the samples was subjected to Sanger sequencing at the Laboratory of Molecular Biology Techniques at the Faculty of Biology, Adam Mickiewicz University in Poznań, Poland.

### 2.7 RNase R treatment

Total RNA extracts from GBM patients’ tissues were treated with RNase R according to the manufacturer’s protocol (Lucigen). Briefly, 2 μg of total RNA was treated with 4 U of RNase R at 37°C for 30 min, followed by 20 min at 65°C. Next, total RNA was purified utilizing NucAway Spin Columns (Invitrogen) and reverse transcribed with the Transcriptor High Fidelity cDNA Synthesis Kit (Roche) according to the manufacturer’s protocol. The PCR was performed – 38 cycles, starting with 3 min of denaturation followed by: 15 sec of denaturation at 95°C, 30 sec of annealing at 63°C, and 20 sec of extension at 72°C. The depletion of linear RNA was confirmed by separating the PCR products in 1.5% agarose gel. Undigested samples served as control.

### 2.8 Subcellular fractionation

Cytoplasmic and nuclear fractions were separated according to the protocol described by Conrad and Ørom. [12] Briefly, a total of 5 × 10^6^ U-138 MG and U-251 MG cells were washed in PBS and incubated with trypsin at 37°C for 5 minutes. The trypsinization was inhibited by adding 10 ml cold EMEM. The cell suspension was transferred into a Falcon tube and centrifuged at 200 x g for 5 minutes. The cell pellet was suspended twice in PBS and centrifuged at 200 x g for 5 minutes. The cell pellet was lysed with Igepal lysis buffer comprised of 10 mM Tris pH 7.4, 150 mM NaCl, 0.15% Igepal CA-360 and 20 U/ml of Protector RNase Inhibitor (Merck) for 5 minutes. The cell lysate was overlaid on top of the sucrose buffer comprised of 10 mM Tris pH 7.4, 150 mM, 24% sucrose, and 20 U/ml of Protector RNase Inhibitor (Merck) and centrifuged at 3500 x g for 10 minutes to obtain separated cytoplasmic and nuclear fractions. The cytoplasmic fraction was centrifuged at 14,000 x g for 1 min and the pellet containing nuclei was rinsed with 1 ml ice-cold PBS-EDTA and centrifuged at 3500 x g for 5 seconds. All centrifugation steps were carried out at 4°C and each incubation was performed on ice. Finally, the RNA was extracted from both fractions using 1 ml of TRIzol reagent per 200 μl of each fraction.

### 2.9 Silencing CLIP2 with siRNA

70-80% confluent cells or spheroids were transfected with siRNAs into a final concentration of 100 nM utilizing siPORT™ Amine as a transfection agent (Thermo Fisher Scientific). Scrambled siRNA was used as a control. Transfected cells were subjected to further experiments 48 h after the transfections. siRNAs (purchased from Sigma-Aldrich) had the following sequences:

1. *non-targeting scrambled siRNA:* 5’-GCUGAACAUGUAGUCACAGAU-3’, 5’-AAGGCACAGCAUGAGCAGGUA-3’;
2. *siRNA circCLIP2:* 5’-AAGGCACAGCAUGAGCAGGUA-3’, 5’ UACCUGCUCAUGCUGUGCCUU-3’;
3. *siRNA mRNA CLIP2:* 5’-GUGUUUGUAACAAUAACGU-3’, 5’-AGUUUAUUGUUACAAACAC-3’.

### 2.10 Proliferation assay

Transfected cells were subjected to proliferation measurement xCELLigence system (Agilent). Cells were seeded onto 96-well Corning Falcon plates in a concentration of 3×10^3^ cells/well in 150 µL of culture media. Proliferation changes were tracked by Adherent-Cell-by-Cell scan type and pictures of proliferating cells were captured every 3 h. The data analysis was performed by software module according to the manufacturer’s recommendations.

### 2.11 Wound healing assay

GBM cells were subjected to wound healing assay to estimate their migratory potential. Briefly U251-MG and U138-MG cell lines were transfected with siRNAs, and 48 hours after, the scratch with a 200 µl pipette tip at the adherent monolayer of the cells was done to generate the wound. The data and images were gathered every 24 hours until the control wound (C) was covered. The data was analyzed by deploying TScratch software.

### 2.12 EMT induction with TGFβ

U-251 MG cells in a concentration of 1×10^5^ cells/mL were seeded onto 6-well plates in a supplemented growth medium and 24 hours after treated with TGFβ to a final concentration of 5 ng/mL. U251-MG-derived spheroids were treated with TGFβ in the same way as adherent cells. Standard conditions for optimal growth and maintenance: 37°C, 95% humidity, and 5% CO_2_ were applied. Next, the cells were harvested at 24 and 48 h time points. Upon the harvest, RNA was extracted and subsequent qRT-PCR performed to check the expression of EMT markers and circRNA.

### 2.13 Invasion assay

The invasion assay was carried out according to the IncuCyte® S3 3D Spheroid Invasion Assay protocol. The invasive potential of U-251 MG- and U-138 MG-derived spheroids was assessed upon the transfection with siRNA. Briefly, spheroids were transfected and incubated for 48 hours with transfection mixture. The Matrigel® (Corning Cat. No. 356234) was diluted into final protein concentration of 2.8 mg/mL and added on the top of the medium. Next, the plate was incubated at 37° C for 30 minutes till the Matrigel® polymerized. Then, the plate was removed from the incubator and 50 µL/well of complete culture media was added gently. The invasion changes were monitored for 4 days.

### 2.14 Magnetic separation of glioma stem cells (GSCs) fraction

U-251 MG- and U-138 MG-derived spheroids were cultured for 2 weeks and subjected to magnetic separation, based on CD133/1 (prominin) antibody-labeled beads recognizing epitope 1 of the CD133 antigen exhibited by the GSCs. Magnetic separation was performed according to the CD133 MicroBead Kit manufacturer’s protocol (Miltenyi Biotec). First, spheroids were gently dissociated with StemPro Accutase (ThermoFisher Scientific). Next, the cells were magnetically labeled and separated using pre-prepared LS columns. Obtained fractions – the population of GSCs (CD133^+^ fraction) and the flowthrough (FT, CD133^−^ fraction) were used for RNA isolation and subsequent qRT-PCR evaluating the expression levels of GSCs markers and circRNAs.

### 2.15 Hypoxia treatment

In order to induce oxygen deficiency, adherent cultures and spheroids were cultured in a condition of 1% oxygen for 5 days. The total RNA from the cells was isolated in Trizol as described previously. The expression of hypoxia markers was assessed by qRT-PCR to confirm the triggering of hypoxic conditions.

### 2.16 Xenograft transplantation of U87-MG silenced cells

U87-MG cells at 70-80% confluence were transfected with either scrambled (control) or circCLIP2 siRNAs into a final concentration of 100 nM utilizing siPORT™ Amine as a transfection agent (Thermo Fisher Scientific) as described in section 2.9. U87-MG cells were implanted into the brains of 3 months old xenograft mice (BALB/c Nude, CAnN.Cg-Foxn1nu/Crl) 24 h upon transfections, the approval from The Local Ethics Council for animal experiments (approval # 38/2019). The cells were dissolved in non-supplemented growth medium and diluted 1:1 with matrigel (Corning). The final volume of 0.25 × 10^6^ cells in 3 µl was injected intracranial per mouse in a speed of 0.5 µl/ min via stereotaxic surgery. In total 9 mice were injected with U87-MG cells: 4 mice (1 male, 3 females) injected with cells transfected by scrambled siRNA as the control group and 5 mice (2 males, 3 females) mice with cells transfected by circCLIP2 siRNA as studied group. The tumors were excised 1 week upon the transplant and snap frozen in liquid nitrogen for further RNA isolation.

### 2.17 Nanostring nCounter analysis and data processing

Total RNA was isolated from frozen tumors excised from mice using Trizol reagent (Invitrogen), treated with DNase I and its quality together with integrity were verified by NanoDrop 2000 spectrophotometer and 1.5% agarose gel electrophoresis as described in section 2.4.

Transcriptomic analysis was done on 100 ng of total RNA per sample by the nCounter SPRINT profiler using a Human Tumor Signaling Panel (catalog # NA-XT-CSO-HTS360-12, nanoString) which measured the expression of 360 genes and 10 reference genes according to the manufacturer’s instructions (nCounter XT Assay User Manual, MAN-10023-11).

The raw data were processed to correct background, normalize the expression and establish the ratio, all performed by dedicated nSolver 4.0 software (nanoString). A defined threshold for background correction was 25. Normalization was performed to the geometric mean of the average of 6 positive controls and 10 housekeeping genes all included in the panel. The fold change (FC) of expression was established by comparison between tumors of mice injected with circCLIP2 siRNA-transfected cells vs. scrambled siRNA-transfected cells. The genes with \**P* ≤ 0.05 were considered significantly upregulated and downregulated.

### 2.18 Western blotting

The protein lysates together with pre-stained protein ladder plus (Thermo Scientific) were suspended in protein sample buffer, boiled for 3 minutes at 95°C and separated in 12% polyacrylamide gel. Separated proteins were transferred to the PVDF membrane by wet blotting and blocked for 1 h at room temperature with 5% skimmed milk in PBS containing 0.05% Tween-20 (PBS-T). Next, the membranes were incubated with primary antibodies diluted 1:1000 (Invitrogen) in 5% skimmed milk overnight at 4°C. After incubation, blots were washed 3 times with PBS-T and incubated for 1 h with anti-mouse or anti-rabbit secondary antibody (Thermo Fisher Scientific) diluted 1:10 000 in 5% skimmed milk. The bands were visualized using West Pico Chemiluminescent Substrate (Thermo Fisher Scientific) in The Syngene G: BOX Chemi XX6 instrument (Synoptic).

### 2.19 Statistical analysis of the results

GraphPad Prism ver. 8 was used to perform statistical analysis on the experimental results. The results are presented as mean value with standard deviation (SD). The differences in mean values between the studied and control samples were evaluated using ANOVA variance extended by Bonferroni post-hoc tests or t-test. Statistically significant results were denoted by the following symbols: ns non-significant, *p ≤ 0.05, **p ≤ 0.01, ***p ≤ 0.001, ****p ≤ 0.0001.

## 3. Results

### 3.1 CircCLIP2 is overexpressed in primary tumor tissues of GBM patients and mainly expressed in the cytoplasm of GBM cell lines

First, in order to identify deregulated levels of circRNAs in GBM, the RNA-seq raw data published by Song et al. (2016) was utilized. [13] The edgeR R package was used to detect the back spliced junction sites. [14] The cutoff criteria for the circRNA identification were as follows: the fold change greater than 2 (FC > 2), and the detection of expression at the level equal or greater than 50% in all analyzed samples. circCLIP2 (hsa_circ_0002755) derived from the parental *CLIP2* gene encoding CAP-GLY domain containing linker protein 2, was found to be substantially overexpressed in the most of analyzed GBM samples. [15, 16] Subsequently, the expression level of circCLIP2 was also confirmed by the analysis deposited on the circBase database. [17]

Next, the expression level of circCLIP2 was examined in tumor tissues from our repository of GBM samples (n=12) by RT-qPCR. The commercially available pools of healthy brain tissues (n=23) served as controls. The expression level of circCLIP2 was found to be 9.85-fold higher in primary GBM tissues compared to healthy brain controls (Fig 1A). Simultaneously, the expression level of the linear transcript of the CLIP2 gene was not significantly elevated (Fig 1A). We found a positive correlation between circCLIP2 expression and the expression of cognate linear mRNA in primary GBM tumor samples. Pearson correlation was equal to 0.8376 (Fig. 1B). This indicates that higher expression of circCLIP2 is assisted with higher expression of its linear transcript. The existence of a circular form of *CLIP2* was confirmed by RNase R treatment while Sanger sequencing confirmed the presence of back-splice site at exon 6 and exon 5 of *CLIP2* gene (Fig. 1C-E).

**Figure 1.**
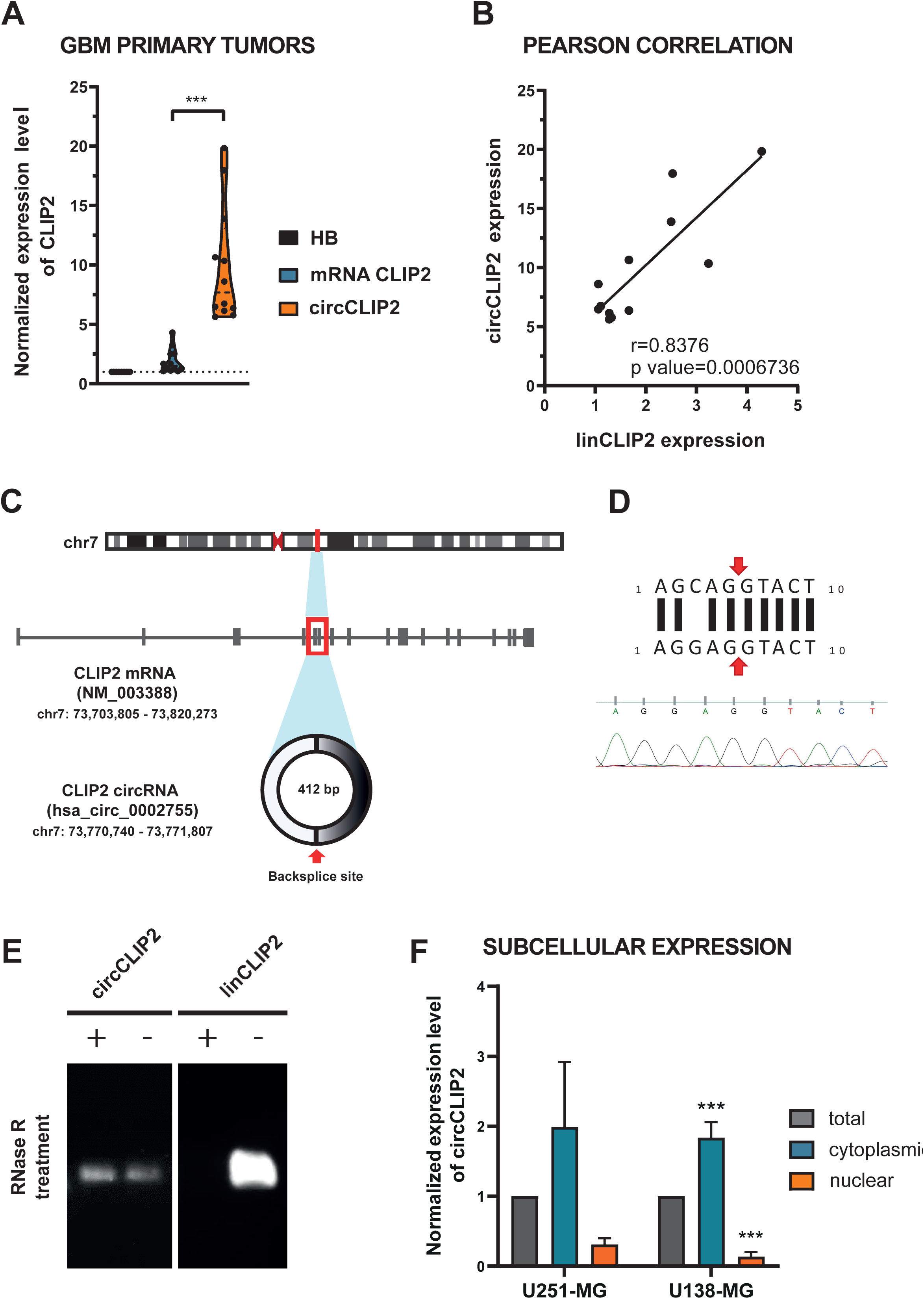
General characteristics and expression levels of linear and circular *CLIP2*. **A** Validation of RNAseq data by qRT-PCR. The expression levels of circCLIP2 and its corresponding mRNA indicate the overexpression of circCLIP2 in primary GBM tumors excised from patients (n=12). The expression level of both transcripts is compared to the control of pooled healthy brains. **B** A positive Pearson’s correlation is calculated for the circCLIP2 and linCLIP2 expression levels in primary GBM tissues. **C** The genomic loci of the *CLIP2* gene and circCLIP2 (hsa_circ_0002755). The red arrow indicates the backsplice site of circCLIP2. **D** Results from Sanger sequencing of the PCR product of circCLIP2 confirm the presence of backsplice junction sequence. **E** Visualization of CLIP2 transcripts upon RNase R treatment. **F** Cytoplasmic localization of circCLIP2 upon subcellular fractionation of U251-MG and U138-MG cells validated by qRT-PCR. \*\*\**P* ≤ 0.001.

To explore the localization and possible mechanism underlying the effects of circCLIP2 in GBM, we performed cellular fractionation. We extracted the total RNA from the nuclear and cytoplasmic fractions of U251-MG and U138-MG cells, respectively. qRT-PCR analysis further clearly showed that circCLIP2 is primarily localized in the cytoplasmic compartment (Fig. 1F, Suppl. Fig. 1).

### 3.2 CircCLIP2 is silenced independently from its linear counterpart in GBM cell lines

Next, in order to find a proper *in vitro* model for further studies, the baseline expression levels of both isoforms of *CLIP2* gene in three adherent cell lines of GBM: U87-MG, U251-MG and U138-MG were established. We observed a significant upregulation of circCLIP2 expression levels in two cell lines U87-MG and U138-MG which were up to 3-fold compared to the pools of healthy brain controls (Fig. 2A).

**Figure 2.**
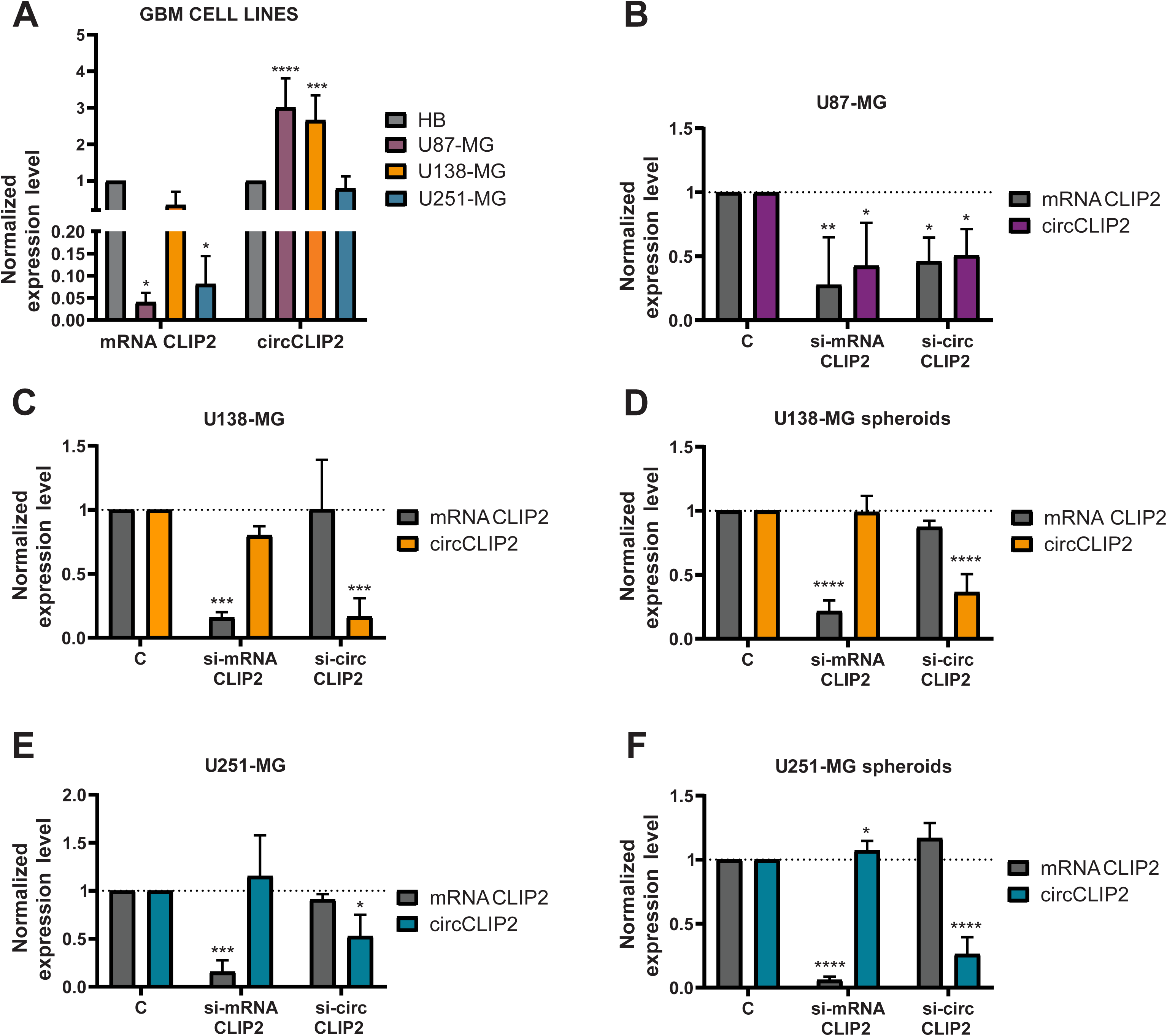
Determination of baseline CLIP2 expression and downregulation specificity upon siRNA silencing in different cell lines evaluated by qRT-PCR. **A** Elevated expression levels of circCLIP2 and its corresponding mRNA present in U87-MG and U138-MG cell lines. The expression level of both transcripts is compared to the control of pooled healthy brains. **B-F** Silencing of linear and circular transcripts of the *CLIP2* gene in adherent U87-MG **(B)**, U138-MG **(C)**, and U251-MG **(E)** cell lines as well as derived spheroids from U138-MG **(D)** and U251-MG **(F)** cell lines. The specificity of knockdown was established by qRT-PCR analysis and expression levels were normalized to scrambled siRNA (C). *p ≤ 0.05, **p ≤ 0.01, ***p ≤ 0.001, ****p ≤ 0.0001.

The effectiveness of siRNA targeting either linear or circular form of CLIP2 in all three adherent cell lines and 3D spheroids derived from U251-MG and U138-MG cell lines was checked. U87-MG cell line purchased from ATCC forms spheroids spontaneously thus no spheroid induction was applied.

We also knockdown the expression level of circCLIP2 using two independent siRNAs (si-mRNA CLIP2 and si-circ CLIP2) to explore the biological function of circCLIP2 and separate it from the function of the cognate mRNA. The knockdown efficiency of siRNAs was determined by qRT-PCR (Fig. 2B-F) compared to the scrambled siRNA control (C) after 48 h upon transfections. The highest silencing efficiency of circCLIP2-nearly 80% knockdown was observed in adherent U138-MG, while U87-MG as well as adherent U251-MG and U251-MG-derived spheroids displayed around 50% drop in the expression of circCLIP2 (Fig. 2B-F). Importantly, transfections of si-circ CLIP2 did not induce significant decrease in the linear form of CLIP2 in the cell lines (Fig. 2B-F), which overall indicates a high specificity of silencing. U251-MG and U138-MG cell lines were selected as *in vitro* models for further functional studies on the role of CLIP2 in GBM while U87-MG cell line was chosen for xenograft transplant due to previously published studies which indicated this cell line as the one forming well-vascularized tumors quickly and recapitulating the GBM biological features when transplanted into xenograft models. [18]

### 3.3 Decreased expression of circCLIP2 diminishes proliferation and migration rates of GBM cells

To initially identify the biological function of circCLIP2 in GBM, we first assessed proliferation of GBM cells. The proliferation rate of the cells upon the circRNAs knockdown was monitored in a real-time mode for 24 to 72 h and measured by xCELLigence system (Agilent). We noticed that silencing of circCLIP2 showed a substantial drop-in proliferation rate: as measured on the about 70% in the tested GBM cells 72 h upon performed transfections (Fig. 3A and C, respectively). Importantly, the silencing of cognate mRNA CLIP2, did not resulted with any changes in proliferation.

**Figure 3.**
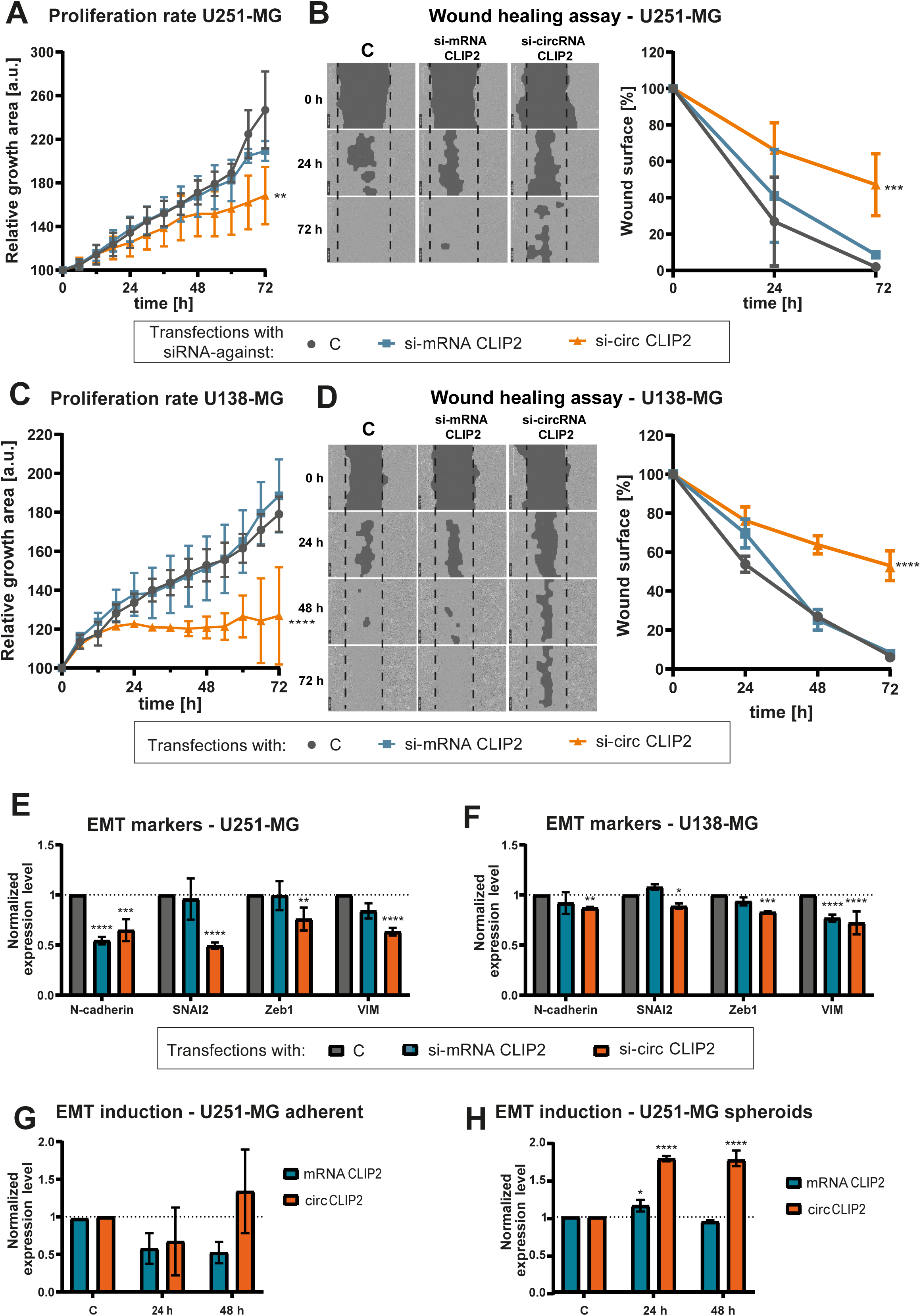
Functional tests assessing proliferation and migration rates upon circCLIP2 knock-down in GBM cells. Determination of EMT-related markers by qRT-PCR. **A** and **C** Proliferation rates were monitored 0-72 h for U251-MG **(A)** and U138-MG **(B)** after circCLIP2 or linear CLIP2 knock-downs. **B** and **D** Wound healing assay was performed for 0-72h in U251-MG (**B**) and U138-MG (**D**) after circCLIP2 or linear CLIP2 knock-downs. Pictures were captured until the wound was covered by cells transfected with scrambled siRNA (control, C). **E-F** Expression levels of EMT markers upon silencing of mRNA CLIP2 or circCLIP2 in U251-MG (**E**) and U138-MG (**F**) cell lines established by qRT-PCR. The expression levels were normalized to scrambled siRNA (control, C). **G**-**H** Analysis of expression levels of CLIP2 transcripts upon EMT induction in U251-MG adherent cell line (**G**) and U251-MG spheroids (**H**). The results were normalized to untreated adherent U251-MG cells and spheroids (control, C). *p ≤ 0.05, **p ≤ 0.01, ***p ≤ 0.001, ****p ≤ 0.0001.

Further, the migratory potential of the GBM cells was explored by utilizing the wound healing assay for 24 to 72 h. The largest area not covered by either U251-MG or U138-MG were upon circCLIP2 knock-down and accounted for 40% and 53% of the initial wound area, respectively (Fig. 3B and D). These results indicate a lower migratory potential of both cell lines expressing diminished levels of circCLIP2 compared to the controls. Similar to proliferation results, none statistically significant effect was observed upon mRNA CLIP2 silencing (Fig. 3B and D).

The results display a potential regulatory role of circCLIP2 in proliferation and migration processes of GBM cells.

### 3.4 Slower migration rate of GBM cells upon circCLIP2 silencing is linked to suppressed EMT. Expression of circCLIP2 is increased in EMT

To determine whether circCLIP2 exerts its tumor-promoting effect by inducing epithelial-to-mesenchymal transition (EMT), we conducted the EMT markers profiling. The expression pattern of EMT markers upon the knock-down of both *CLIP2* gene isoforms in U251-MG and U138-MG cell lines was analyzed by qRT-PCR. The expression of all analyzed EMT markers, namely N-cadherin, SNAI2, Zeb1 and vimentin (VIM) was significantly decreased upon circCLIP2 knock-down in both cell lines (Fig. 3E-F). The most significant drop, namely 51% and 37%, was observed in the expression of SNAI2 and VIM in U251-MG cell line, respectively. Interestingly, silencing of linear CLIP2 affected the expression of N-cadherin in U251-MG and VIM in U138-MG cell lines (Fig. 3E-F).

Next, we inversely induced EMT process to better illustrate the tumor-promoting effects by circCLIP2 via EMT in GBM. Thus, adherent U251-MG cell line and U251-MG spheroids were treated with TGFβ for 24 and 48 h to induce the EMT. Untreated U251-MG adherent cells and spheroids served as controls (C). The conformation of successful EMT induction was evaluated by measuring the expression levels of EMT markers by qRT-PCR. The highest upregulation in expression level was observed for SNAI2: 6.43-fold and 4.47-fold change in adherent U251-MG cells and spheroids, respectively (Suppl. Fig. 2A-B). Further, the expression level of both circular and linear *CLIP2* isoforms was validated under the condition of induced EMT (Fig. 3G-H). The elevated expression of circCLIP2 was observed as ∼1.8-fold change only in U251-MG spheroids 24 and 48 h after the TGFβ stimulation (Fig. 3G-H). Altogether, these results point out that circCLIP2 is upregulated during EMT and promotes migration processes.

### 3.5 circCLIP2 expression is specific for GSC and elevated under hypoxia. Downregulated expression of circCLIP2 results in diminished stemness and hinders invasion

Since GSCs are key drivers of tumor growth and resistance to therapy, thus we also aimed to study their relation to circCLIP2. Since deprivation of oxygen is known to maintain the stemness of GBM tumors, the hypoxic conditions were induced in GSC-rich spheroids derived from U251-MG and U138-MG cell lines. First, a successful induction of hypoxia conditions was verified by the evaluation of GLUT1, ANG and PDK1 expression levels by qRT-PCR and western blotting (Suppl. Fig. 3A-C). A substantial increase in the expression of those hypoxia activated genes in both U251-MG and U138-MG spheroids was revealed based on qRT-PCR analysis. U251-MG spheroids displayed a stronger response to oxygen deficiency since expression of hypoxia markers was higher (7.5-fold, 8.62-fold, and 6.27-fold change for GLUT1, ANG and PDK1, respectively), in comparison to U138-MG-derived spheroids (3.55-fold, 2.43-fold and 2.52-fold change for GLUT1, ANG, and PDK1, respectively) which was confirmed by qRT-PCR (Suppl. Fig. 3A). The western blot analysis confirmed elevated levels of GLUT1 and PDK1 at the protein level of both U251-MG and U138-MG cell lines (Suppl. Fig. 3B-C). To address whether hypoxic conditions influence the expression of *CLIP2* isoforms, the spheroids were cultured in hypoxia conditions for 5 days, harvested and assayed by qRT-PCR (Fig. 4A). CircCLIP2 expression level was found elevated in both types of spheroids: the increase of 3.38-fold change for U251-MG-derived spheroids and the increase of 1.58-fold change for U138-MG derived-spheroids, respectively. While the expression level of linear *CLIP2* was significantly elevated by 2.07-fold exclusively in U251-MG-derived spheroids (Fig. 4A).

**Figure 4.**
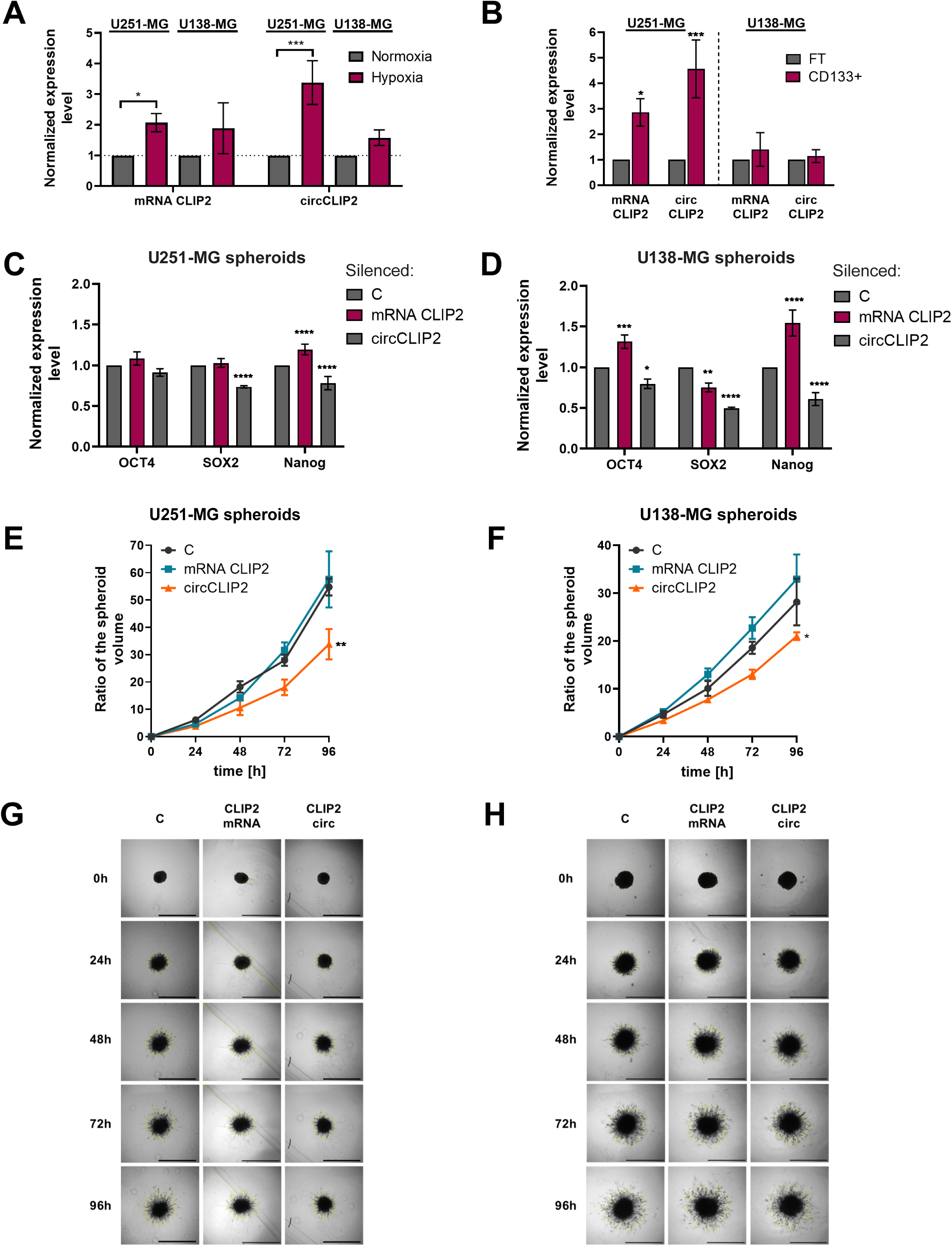
Expression of CLIP2 under hypoxia conditions. Expression of pluripotency markers upon CLIP2 knock-downs. Invasion rates of spheroids with silenced CLIP2. **A** Expression levels of CLIP2 isoforms in GBM cell lines under hypoxia conditions established by qRT-PCR. The results were normalized to cells grown under normoxia conditions. **B** Expression levels of CLIP2 in CD133 positive (CD133^+^) and negative (flow-through, FT) fractions isolated from U251-MG spheroids and U138-MG spheroids. **C-D** Expression level of GSC markers in spheroids derived from U251-MG (**C**) and U138-MG (**D**) upon CLIP2 silencing. **E-H.** Volume ratio of spheroids derived from U251-MG (**E**) and U138-MG (**F**) grown for 96h upon CLIP2 silencing. Representative microscopy pictures of U251-MG (**G**) and U138-MG (**H**) spheroids taken every 24 h. The results of functional assay were normalized to cells transfected with scrambled siRNA (control, C). *p ≤ 0.05, **p ≤ 0.01, ***p ≤ 0.001, ****p ≤ 0.0001.

In order to study the potential role of circCLIP2 in GSCs, their fractions were enriched and separated from U251-MG- and U138-MG-derived spheroids. The fraction compared to their adherent counterparts display higher protein levels of stemness markers as indicated by western blotting (Suppl. Fig. 4A). Magnetic separation was done with antibody against CD133^+^ (prominin), a specific surface marker expressed by GSCs. The purity of fractions: stem cell-rich (CD133^+^) compared to non-GSCs fractions (remained in the flow-through, FT) was validated by the expression analysis of the pluripotency markers *OCT4*, *SOX2* and *Nanog* by qRT-PCR. Higher expression of *OCT4* and *Nanog* was observed in CD133^+^ fraction compared to FT fraction in U251-MG-derived cell lines while only the expression of *OCT4* was significantly elevated in CD133^+^ fraction from U138-MG-derived cell lines (Suppl. Fig. 4B-C). When the purity of fractions was confirmed, the expression of mRNA CLIP2 and circCLIP2 was analyzed by qRT-PCR. Both mRNA CLIP2 and circCLIP2 were found highly expressed in CD133^+^ of U251-MG-derived spheroids (2.84-fold change and 4.43-fold change, respectively) normalized to FT, while no elevation in expression of any isoform of *CLIP2* was found in U138-MG-derived spheroids (Fig. 4B).

Next, to verify whether oxygen deprivation could additionally enhance the expression of circCLIP2, the CD133^+^ fraction was separated from U87-MG cells grown under hypoxia conditions for 5 days. Substantial increase in expression of solely circCLIP2 isoform in CD133^+^ fractions was observed under both normoxia and hypoxia conditions. Interestingly, the expression of circCLIP2 doubled in CD133^+^ fraction grown under hypoxia compared to the fraction of CD133^+^ cells grown under standard oxygen conditions (Suppl. Fig. 4D).

To check whether circCLIP2 could impact the stemness of GBM cells, the qRT-PCR analysis was performed on U251-MG- and U138-MG-derived spheroids upon siRNA silencing of circCLIP2 (Fig. 4C-D). U251-MG-derived spheroids with downregulated levels of circCLIP2 displayed a significant drop in the expression levels of *SOX2* (0.74-fold change) and Nanog (0.78-fold change) markers (Fig. 4C) while U138-MG-derived spheroids showed a decrease in the expression of *OCT4* (0.8-fold change), *SOX2* (0.5-fold change), and *Nanog* (0.6-fold change) markers (Fig. 4D). Interestingly, the knock-down of linear CLIP2 transcript led to statistically significant increase in the levels of stem cell markers especially in U138-MG-derived cell line (Fig. 4C-D). These results clearly show that circCLIP2 plays a regulatory role specifically in stem cell fraction of GBM and its expression gets elevated under hypoxia condition.

Further, the functional test was performed to study the invasive properties of spheroids derived from both cell lines upon circCLIP2 silencing monitored for 96 h (Fig. 4E-H). Silencing of circCLIP2 in both U251-MG- and U138-MG-derived spheroids resulted in aa slower rate of invasion compared to spheroids transfected with scrambled siRNA. The invasion rate was established based on calculating the volume of spheres (Fig. 4E-H). A significant drop of invasion rate was observed for U251-MG- and U138-MG-derived spheroids, 0.62-fold change and 0.75-fold, respectively (Fig. 4E-H). The knock-down of the mRNA CLIP2 did not impact the invasion rate of any spheroids utilized since it was comparable with control – spheroids transfected with scrambled siRNA (Fig. 4E-H). These results highlight the role of circCLIP2 in invasion process of not only adherent GBM cells but also in GSC-rich 3D structures.

### 3.6 circCLIP2 silencing mediates inhibition of xenograft tumor growth via EMT and ECM

Next, we sought to confirm the tumor-promoting effect of circCLIP2 *in vivo* using the xenograft model of brain tumors. U87-MG cells were transfected with scrambled (control) or circCLIP2-siRNAs and the silencing effect was monitored for 3 days. The silencing effect of circCLIP2 siRNA was observed to be the highest 72 h upon transfection - 25% drop in the expression of circCLIP2 compared to transfection with scrambled siRNA (Fig. 5A-C). The silenced cells was collected 24 h after transfection and intracranially injected into the mice. The experiment was terminated 1 week after performed transplant. The results revealed a trend of a decrease in tumor volume as calculated by ImageJ program (Fig. 5D). Based on qRT-PCR analysis, the effect of circCLIP2 silencing was no longer observed in tumor mass 1 week upon the transplant and excision as expected since siRNA-mediated knock downs were of a transient type (Fig. 5E).

**Figure 5.**
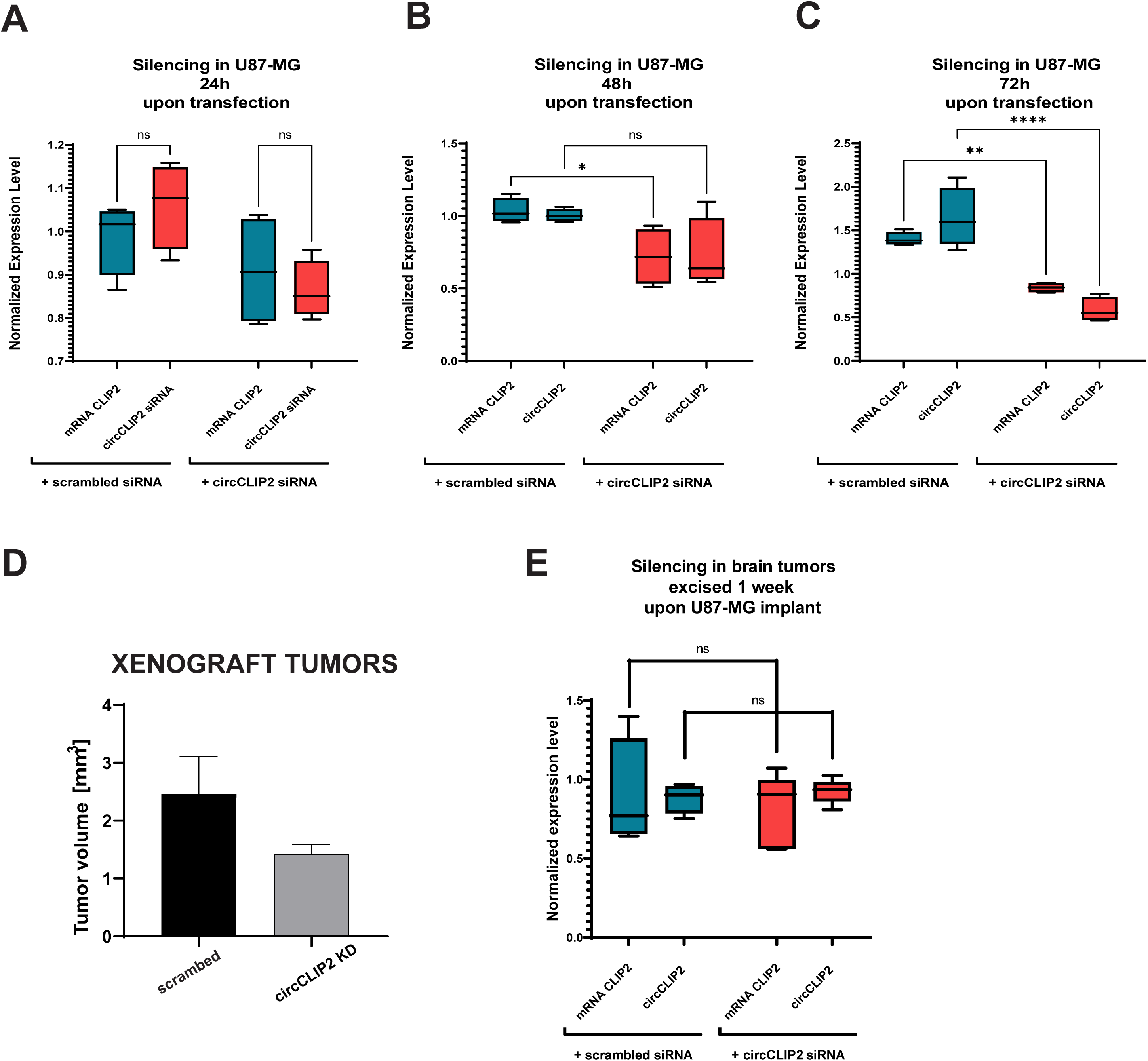
Expression of CLIP2 upon silencing of U87-MG cells prior xenograft transplant. The volume of tumors excised 1 week after xenograft transplant and the silencing effect. **A-C** Transfection efficiency in U87-MG cells was monitored every 24 h for 72h. A pool of transfected cells was injected intracranially 24 h upon cell transfection with either scrambled or circCLIP2-targeting siRNA. **D** The volume of tumors (mm^3^) was calculated by ImageJ program. **E** No silencing effect of circCLIP2 is observed in excised tumors 1 week after the transplant as evaluated by qRT-PCR. ns non-significant, *p ≤ 0.05, **p ≤ 0.01, ****p ≤ 0.0001.

To further explore the molecular changes related to the physiological observations upon circCLIP2 knockdown, we isolated RNA from resected tumors. Then, the expression profiling by the panel of tumor signaling pathway (Nanostring) and qRT-PCR analysis was performed. A total of 44 genes were found to be differentially expressed from the panel list (statistical significance p < 0.05). The cut off of minimum 0.2 increase or decrease in fold change expression was chosen. Thus, finally 22 downregulated and 22 upregulated genes in siRNA-circCLIP2-silenced tumors compared to scrambled siRNA. The changes in expression of distinct genes presents the heatmap (Fig. 6A). The expression of master modulator of ECM composition, *MMP9* was found downregulated together with ECM-binding and signaling receptors, integrins *ITGA5*, *ITGB3* and *ITGA11* which expression was also significantly changed. Interestingly, the expression of both receptor NOTCH3 and its ligand JAG1, the components of Notch signaling which facilitates cell-to-cell communication was also decreased.

**Figure 6.**
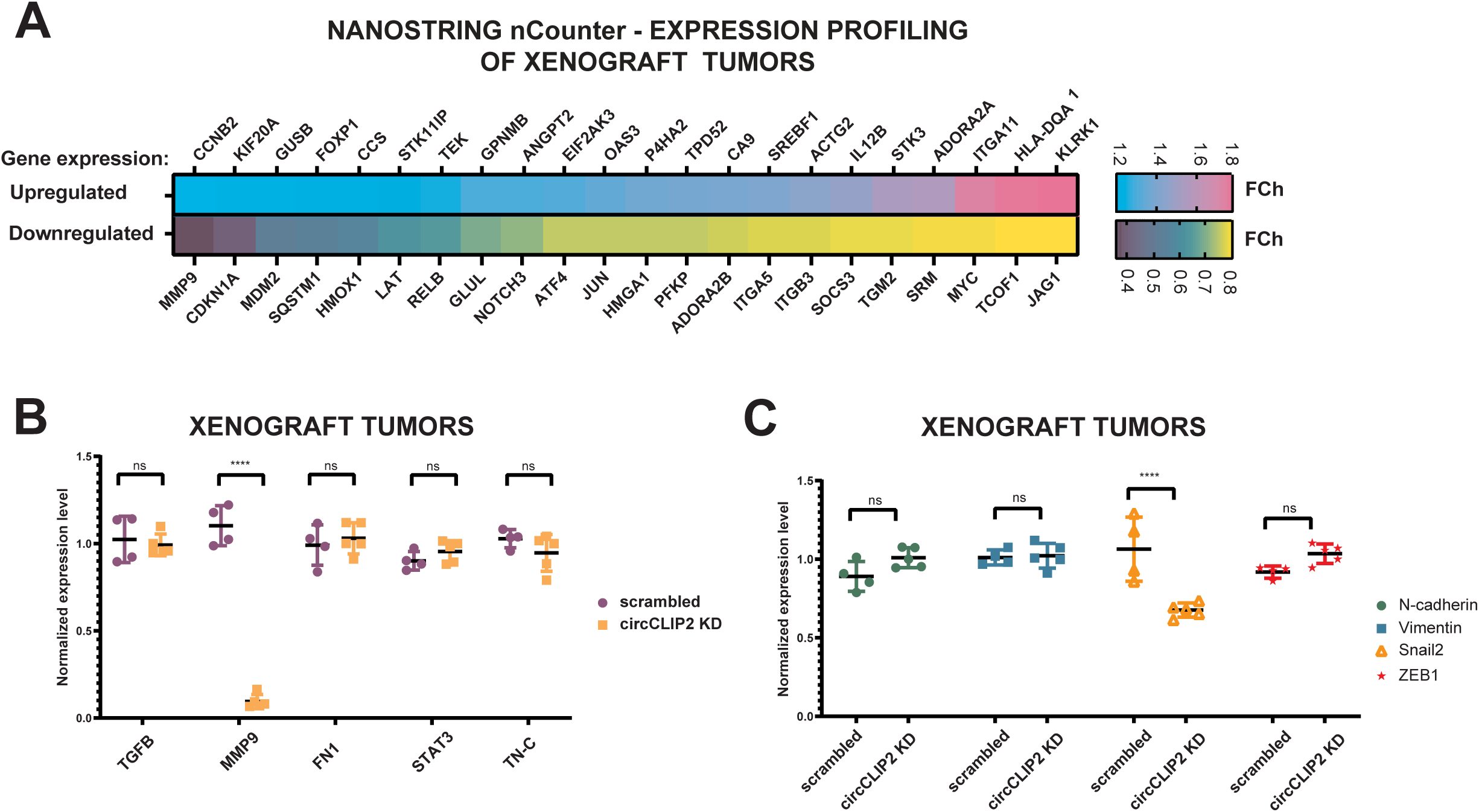
Expression profiling of cancer signaling including ECM and EMT in xenograft tumors. **A** The heatmap was generated based on the fold changes calculated in circCLIP2-silenced tumors and compared to tumors transfected with scrambled siRNA. Genes were selected based on statistically significant (\**P* ≤ 0.05) fold changes between the groups with 0.2 and -0.2 cut-offs. Expression levels are indicated by colored bars from purple (low expression) to pink (high expression). Profiling was done by Nanostring nCounter analysis. **B-C** The expression levels of genes involved in ECM reorganization and signaling (**B**) and genes involved in EMT process were evaluated by qRT-PCR. ns non-significant, ****p ≤ 0.0001.

Although, due to the transient type of siRNA transfections, the effect of circCLIP2 silencing was no longer observed in tumor mass 1 week upon the transplant and excision (Suppl. Fig. 5), the downstream progressive molecular changes were already initiated and maintained. Among the predominant changes observed was the profound downregulation of the metalloproteinase 9 (*MMP9*) (Fig. 6B). Its expression was validated in 1-week xenograft tumors by qRT-PCR, together with the expression of genes affecting extracellular matrix (ECM) composition and interactions: *TGFB*, *FN1*, *STAT3,* and *TNC* during carcinogenesis as confirmed by previous experimental studies.

Additional qRT-PCR analysis did also show the significant changes in the expression of EMT-related gene *Snail2* which was found to be downregulated in 40%. The expression of *N-cadherin*, *vimentin* or *ZEB1* was not significantly changed (Fig. 6C).

## 4. Discussion

GBM is the most aggressive and the deadliest type among brain tumors which affects adult population. [19] In addition to malignancy, hard location and high heterogeneity of GBM tumors make the prognosis and treatment extremely challenging. [20–23] In order to find relevant stratification and prognostic markers as well as druggable targets, extensive studies are carried out at the cellular and molecular levels. [24–26] circRNAs belong to a group of promising molecular candidates due to their brain-specific expression. [27–29] However, the exact molecular roles of most circRNAs remain elusive and a strong urge to focus on the role of single candidates identified by global screens exists. [30, 31] Thus, the aim of herein study was to reveal the biological role of circCLIP2 in GBM found highly expressed in the tumor tissues constituting in established repository. Elevated expression of circCLIP2 in patient samples was confirmed and additionally found in GBM cell lines and derived spheroids, specifically in GSC fraction under hypoxia. A significant knockdown of circCLIP2 expression was also established independently from its linear counterpart which enabled further functional studies. Lower rates of proliferation, invasion and migration observed upon circCLIP2 silencing indicated this circular RNA as a substantial modulator of those cancer processes and thus, a strong contributor into the aggressiveness of GBM. Profiling of gene expression related to tumor signaling pathways in xenograft tumors upon circCLIP2 silencing, identified deregulated expression of molecules mainly linked with ECM. The expression of the master modulator of ECM composition, *MMP9* was found downregulated together with ECM-binding and signaling receptors - integrins *ITGA5*, *ITGB3* and *ITGA11* which expression was also significantly changed. [32–34] Interestingly, also the expression of both receptor *NOTCH3* and its ligand *JAG1*, which facilitate cell-to-cell communication and determine the cell fate was also decreased assuming the switch in cell fate upon circCLIP2 downregulation. [35] These progressive changes were observed despite the fact, that due to the transient silencing, the circCLIP2 knockdown was no longer observed in tumor mass 1 week upon the transplant and excision.

CircCLIP2 has been previously indicated as an upregulated circular RNA in GBM [13] and as an eventual marker of high-grade serous ovarian cancer (HGSOC) [36] In addition, a few reports display a high upregulation of circCLIP2 in primary brain tumors. [13, 15, 16]

Results presented here confirmed the elevated expression levels of circCLIP2 not only in the tissues of GBM tumors but also in commercially available GBM cell lines. Altogether, the upregulated basal expression of circCLIP2 points toward potential oncogenic function in the biology of GBM. Importantly, an isoform-specific downregulation in the expression of either circCLIP2 or mRNA CLIP2 was obtained, independently by applying two different targeting siRNAs. By gaining such silencing specificity, it was possible to study the exclusive biological role of circCLIP2. Moreover, a substantial decrease of circCLIP2 expression in *in vitro* models enabled for functional studies and xenograft transplant.

siRNA-mediated knock-downs of circCLIP2 led to a drop in proliferative potential of the GBM cells. So far, an impaired cell proliferation upon circCLIP2 knock-down has been reported to occur via miR-195-5p/HMGB3 axis and further activation of Wnt/β-catenin signaling in glioma cells. [16] However, the gene expression profiling in our studies does not indicate any significant deregulation in Wnt signaling to be linked to GBM proliferation but rather a substantial drop in the expression of *CDKN1A*, *ITGB3* and *MYC* genes which together are the positive regulators of proliferation. [37]

The results also demonstrated, that the knock-down of circCLIP2 leads to diminished both GBM cell migration and invasive potential, the processes stimulated via EMT in GBM [38, 39] EMT is a biological process that allows immobile epithelial cells to gain a mesenchymal phenotype and thus plasticity essential for relocation. [40, 41] As EMT increases the migratory, invasive, and metastatic potential, the expression level of circCLIP2 was investigated under EMT-induced conditions applied into both adherent cells and spheroids. The data obtained indicated elevated expression levels of circCLIP2 in 2D and 3D cultures, which shows that EMT induces circCLIP2 expression that likely fuels the aggressiveness of GBM. The decrease in migration and invasion upon circCLIP2 downregulation confirmed this result.

The EMT initiation and progression is obviously effected by tumor microenvironment, including hypoxic niche. [42] Hypoxia was shown to enhance the migration and invasion of GBM by promoting the mesenchymal shift via HIF1α–ZEB1 axis. [43] Our studies also demonstrated that the expression of both isoforms of *CLIP2* gene are upregulated under oxygen deficiency conditions, with a strong prevalence of circCLIP2. The results also show the overexpression of circCLIP2 isoform in the GSCs fraction separated from GBM spheroids. Interestingly, previously published studies indicated that the knock-down of *SOX2*, one of key stemness markers in GSC, led to decreased expression of mRNA CLIP2 in GBM. [44] As our studies show changes in invasion rate of spheroids upon the knock-down of circCLIP2 and changes in EMT markers expression. However, deregulation of EMT markers are not observed in xenograft tumors except for Snail2, which might be explained by too short period of tumor growth upon transplant where just the initial changes could have been scoped including the changes in ECM signaling.

In summary, this study identifies circCLIP2 as novel circRNA associated with GSC, EMT and ECM. Obtained results clearly show the involvement of circCLIP2 in GBM aggressiveness and unravel a part of probably more complex mechanism behind the circCLIP2 action through MMP-9, integrins and Snail2 molecules. Our results open new possibilities for further studies focused on revealing detailed mechanisms which are engaged by circCLIP2 and may be of prognostic and therapeutic value.

## 6. Authors’ Contribution

Conceptualization: JM, ZZ, KK, JLŁ, AG, PG, MPS, and KR; Sample collection: ZZ, JM, KK, JLŁ, AG, PG, AMB, AK, DW; Formal analysis: ZZ, JM, KK, JLŁ, AG, PG, DW, MPS, and KR; Investigation: JM, ZZ, KK, JLŁ, AG, MPS, and KR; Methodology: ZZ, JM, KK, JLŁ, AG, PG, AK, DW, ŁP and KR; Resources: KR; ŁP; Supervision: KR, ŁP; Funding acquisition: KR; Visualization: JM, KK, ZZ, JLŁ, AG; Writing - original draft: ZZ, JM, KK, JLŁ, AG; Writing - review & editing: JM, ZZ, KK, JLŁ, AG and KR.

## 8. List of Abbreviations

GBM: Glioblastoma Multiforme
IDH1: Isocitrate Dehydrogenase 1
IDH2: Isocitrate Dehydrogenase 2
circRNA: circular RNA
HGSOC: High-Grade Serous Ovarian Carcinoma
MAPK: Mitogen-Activated Protein Kinase
TGFβ: Tumor Growth Factor β
GLUT1: Glucose Transporter 1
ANG: Angiogenin
PDK1: 3-phosphoinositide-dependent kinase 1
SNAI1: Snail Family Transcriptional Repressor 1
SNAI2: Snail Family Transcriptional Repressor 2
VIM: Vimentin
GSCs: Glioma Stem Cells
SOX2: Sex Determining Region Y-box 2
EMT: Epithelial – Mesenchymal Transition
TMZ: Temozolomide

## 9. Acknowledgements

We would like to acknowledge: Agnieszka Rybak-Wolf and Nikolaus Rajewsky’s laboratory for sparing nCounter SPRINT profiler, Magdalena Woźna-Wysocka for help in expression analysis based on nSolver software and Małgorzata Grabowska for initial proofreading of the manuscript and insightful comments.

## 10. Ethics approval and consent to participate

The GBM samples (*n* = 12) were obtained from the Clinic of Neurosurgery and Neurotraumatology, Karol Marcinkowski University of Medical Sciences in Poznan, Poland, based on the approval by the Bioethics Council of the Poznan University of Medical Science (Nr. 46/13) and the informed consent obtained from the patients.

## 11. Data Availability Statement

To identify the circRNAs important for the pathological process of GBM, the publicly available data by Song et al. (2016) were used. [13]

---

**Supplemental Figure 1.**
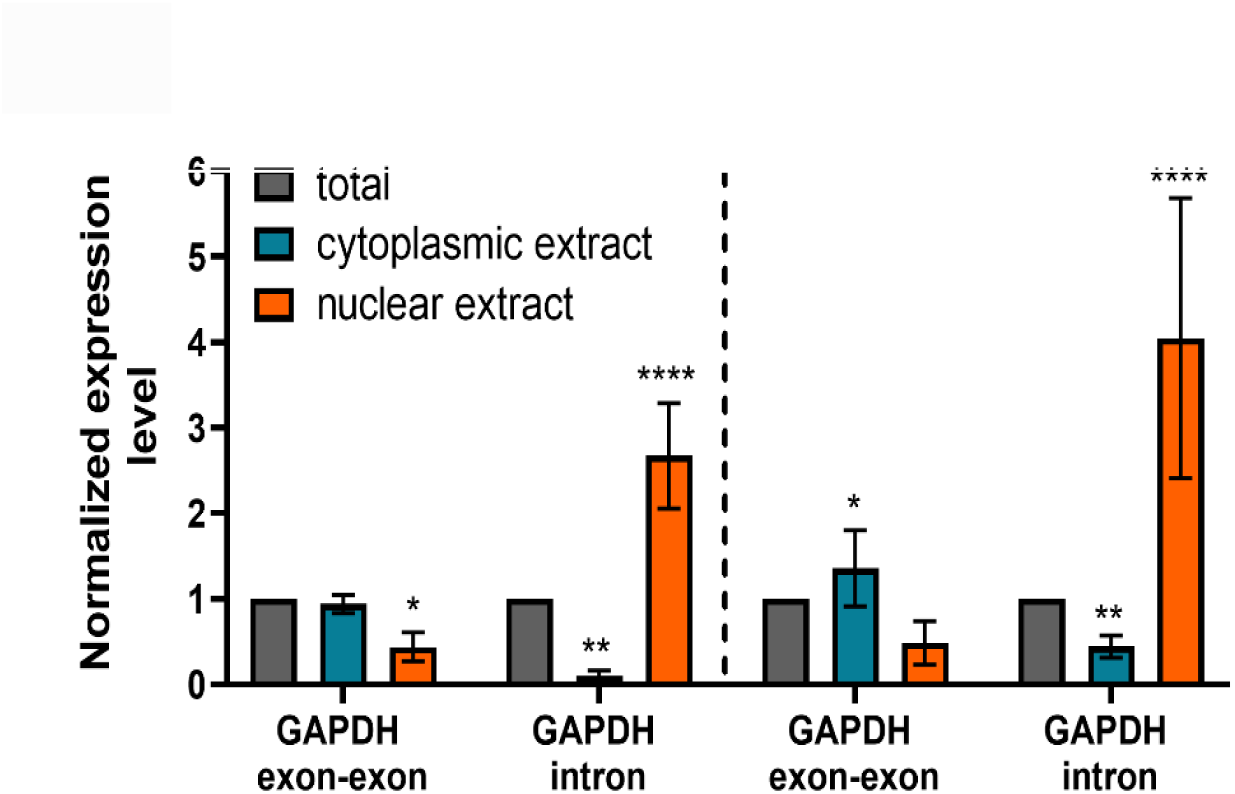
GAPDH subcellular localization upon fractionation. Upon subcellular fractionation of U251-MG (left panel) and U138-MG (right panel) cell lysates, the levels of GAPDH expression were validated by qRT-PCR. *p ≤ 0.05, **p ≤ 0.01, ****p ≤ 0.0001.

**Supplemental Figure 2.**
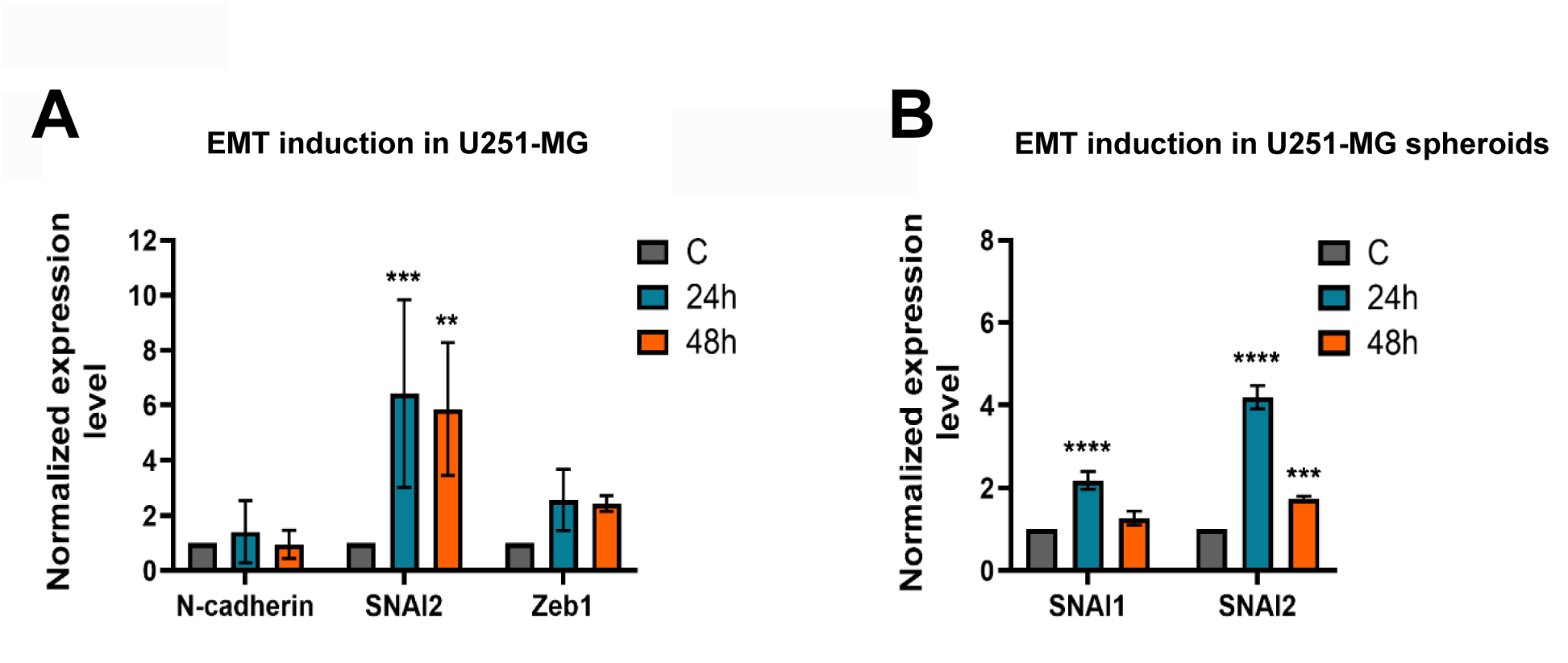
EMT induction by TGFß in U251-MG adherent cells and U251-MG spheroids. **A-B** The induction was verified by measuring the expression of EMT markers upon 24 and 48 h using qRT-PCR. **p ≤ 0.01, ***p ≤ 0.001, ****p ≤ 0.0001.

**Supplemental Figure 3.**
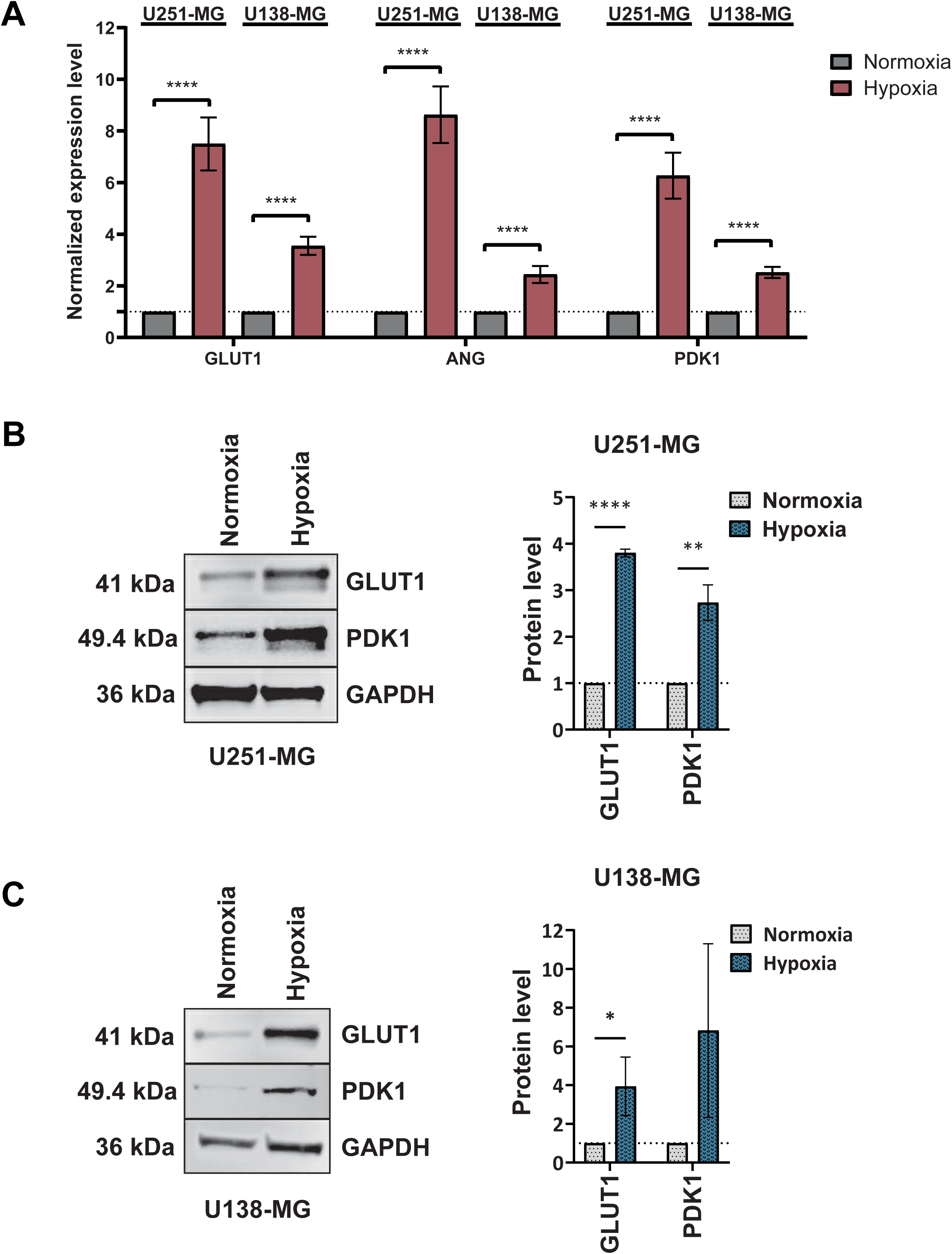
Hypoxia induction in U251-MG and U138-MG adherent cells. **A** The induction of hypoxia was verified by measuring the expression of hypoxia markers upon 48 h using qRT-PCR. **B-C** The induction of hypoxia was verified by measuring the protein levels of hypoxia markers upon 48 h by western blotting. Quantitation of immunoblotting results was done by ImageJ software. *p ≤ 0.05, **p ≤ 0.01, ****p ≤ 0.0001.

**Supplemental Figure 4.**
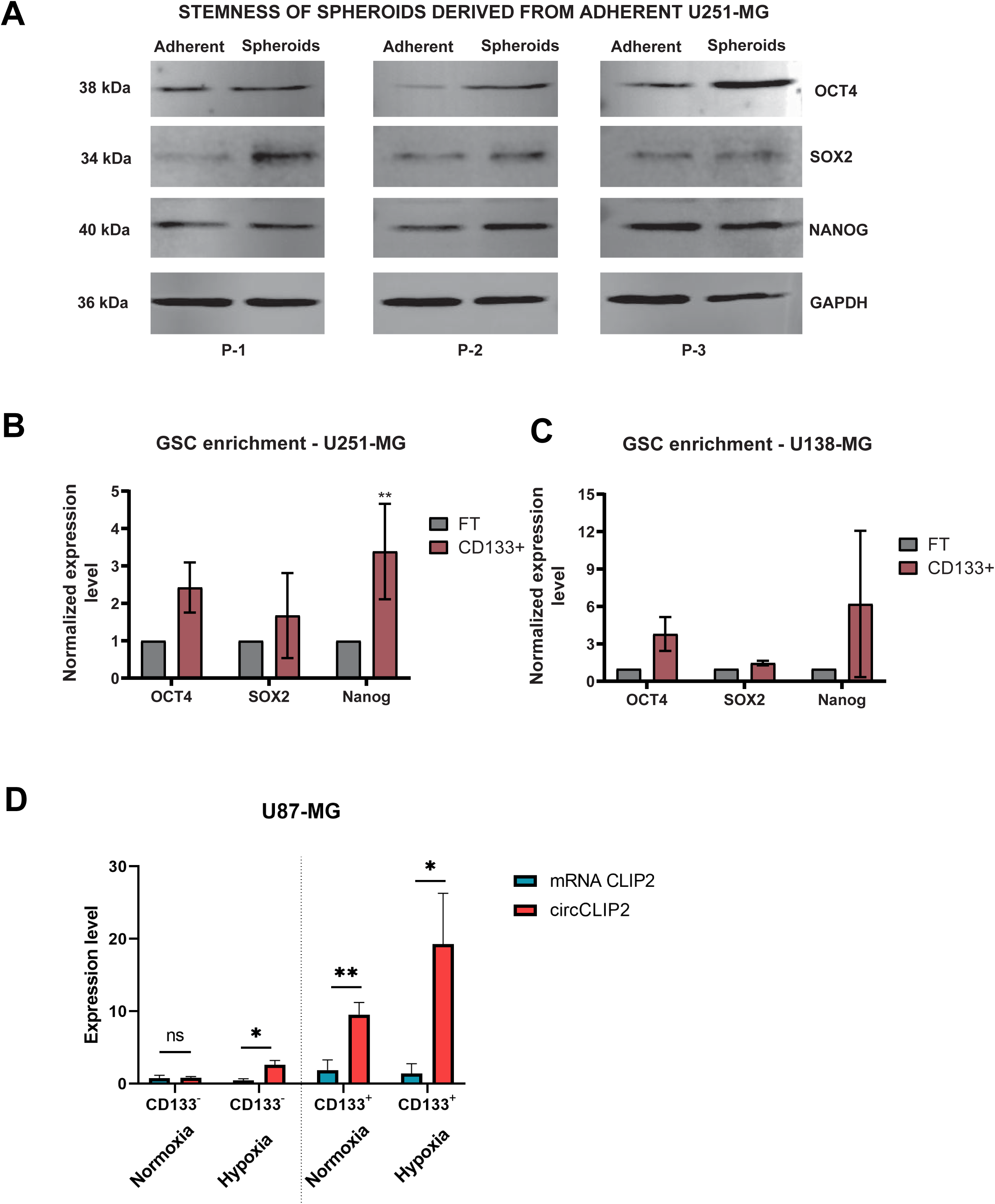
Stemness evaluation under baseline and hypoxia conditions. **A** The stemness of U251-MG was verified by measuring the protein levels of pluripotency markers in both adherent cells and spheroids using western blotting. **B-C** Control experiments in which the expression of pluripotency markers was verified in U251-MG- (**B**) and U138-MG- (**C**) derived spheres upon isolating the CD133^+^ fraction rich in GSCs. The analysis was done by qRT-PCR. **D** *CLIP2* expression shown under normoxia and hypoxia conditions upon CD133^+^ isolation. Expression evaluation was done by qRT-PCR. *p ≤ 0.05, **p ≤ 0.01.

**Supplemental Table 1.**
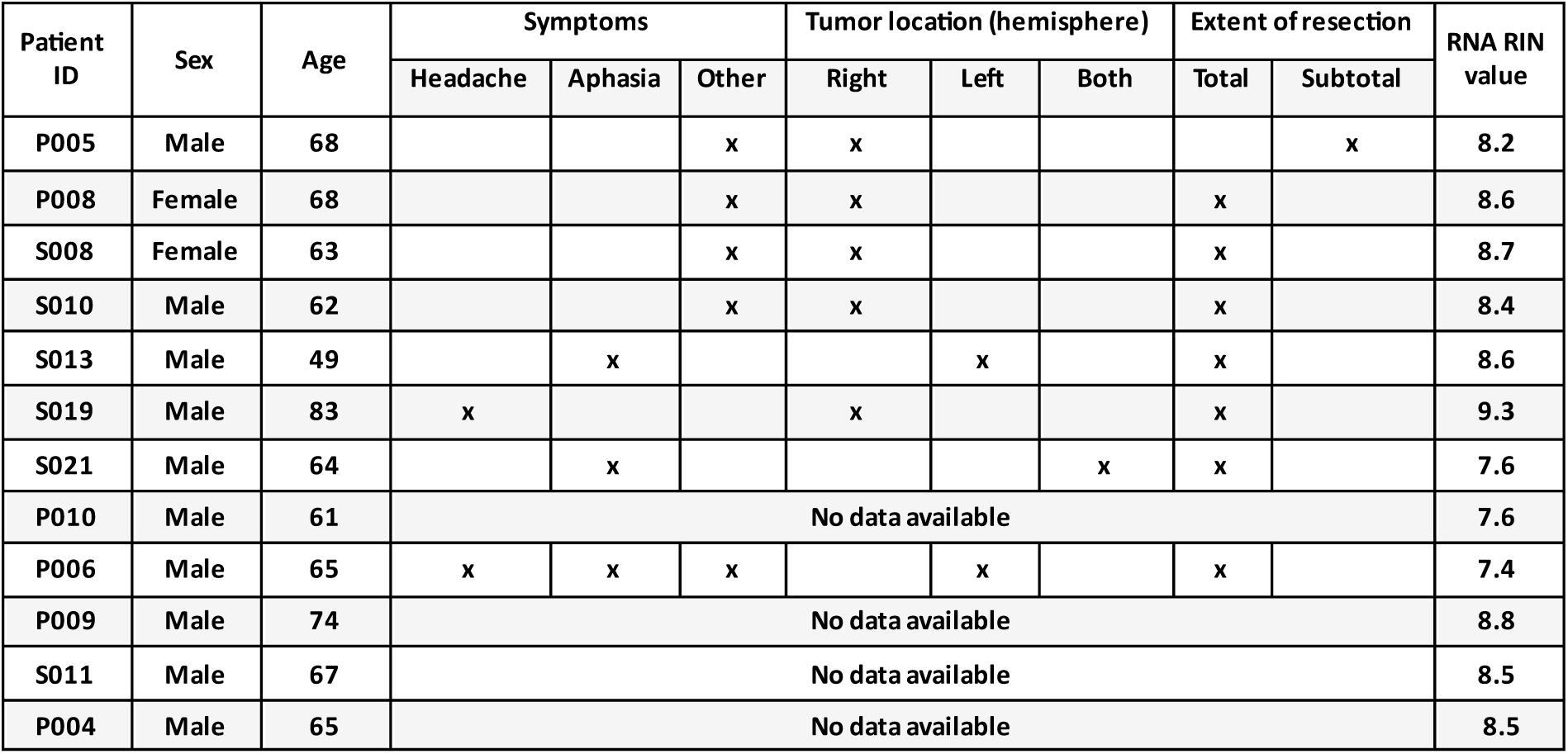
Characterization of GBM patients and tissues used in the studies.

| Patient ID | Sex | Age | Symptoms |  |  | Tumor location (hemisphere) |  |  | Extent of resection |  | RNA RIN value |
| --- | --- | --- | --- | --- | --- | --- | --- | --- | --- | --- | --- |
|  |  |  | Headache | Aphasia | Other | Right | Left | Both | Total | Subtotal |  |
| P005 | Male | 68 |  |  | x | x |  |  |  | x | 8.2 |
| P008 | Female | 68 |  |  | x | x |  |  | x |  | 8.6 |
| S008 | Female | 63 |  |  | x | x |  |  | x |  | 8.7 |
| S010 | Male | 62 |  |  | x | x |  |  | x |  | 8.4 |
| S013 | Male | 49 |  | x |  |  | x |  | x |  | 8.6 |
| S019 | Male | 83 | x |  |  | x |  |  | x |  | 9.3 |
| S021 | Male | 64 |  | x |  |  |  | x | x |  | 7.6 |
| P010 | Male | 61 | No data available |  |  |  |  |  |  |  | 7.6 |
| P006 | Male | 65 | x | x | x |  | x |  | x |  | 7.4 |
| P009 | Male | 74 | No data available |  |  |  |  |  |  |  | 8.8 |
| S011 | Male | 67 | No data available |  |  |  |  |  |  |  | 8.5 |
| P004 | Male | 65 | No data available |  |  |  |  |  |  |  | 8.5 |

**Supplemental Table 2.**
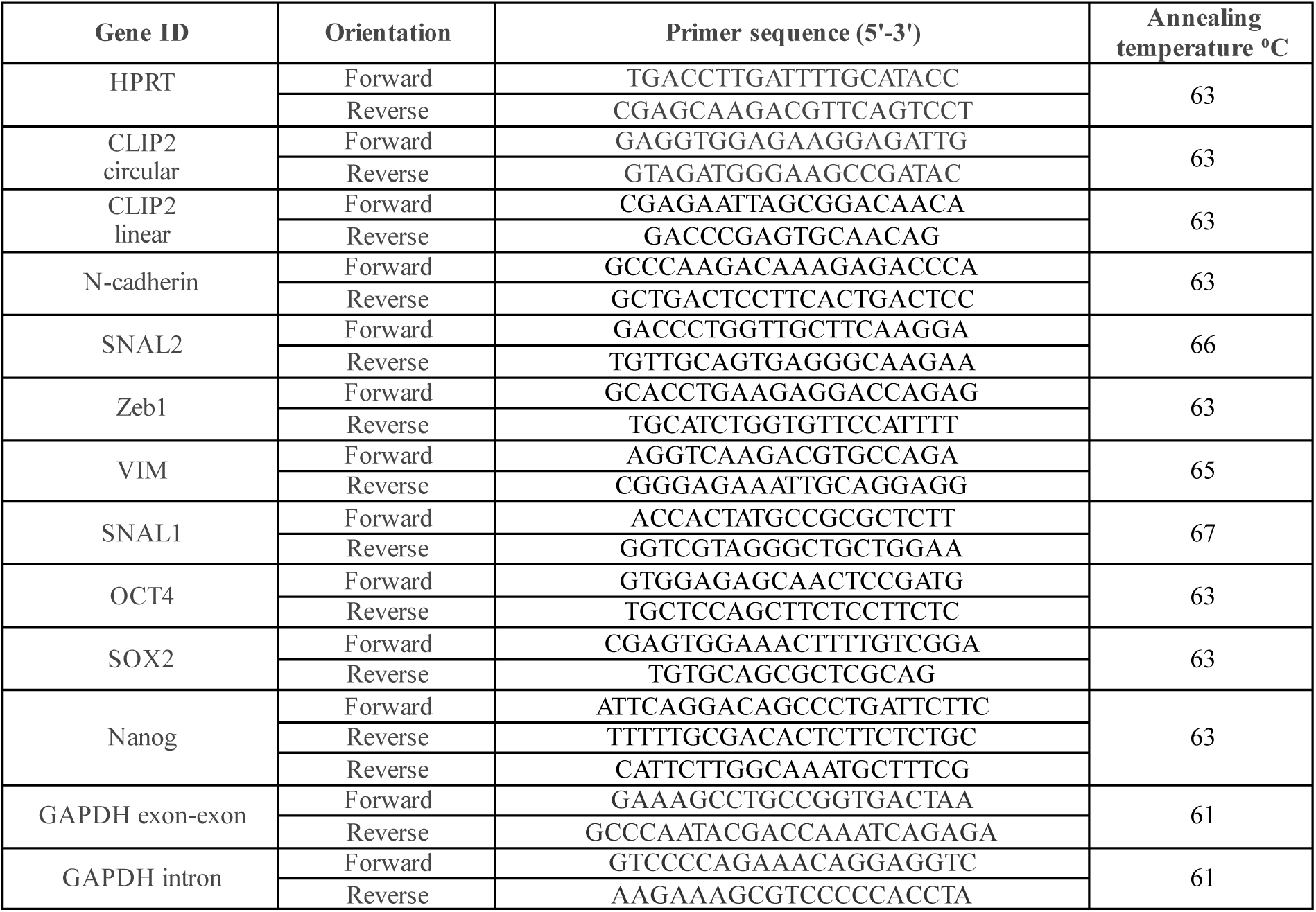
Sequences of primers used for qRT-PCR analyzes.

**Supplementary Table 3.**
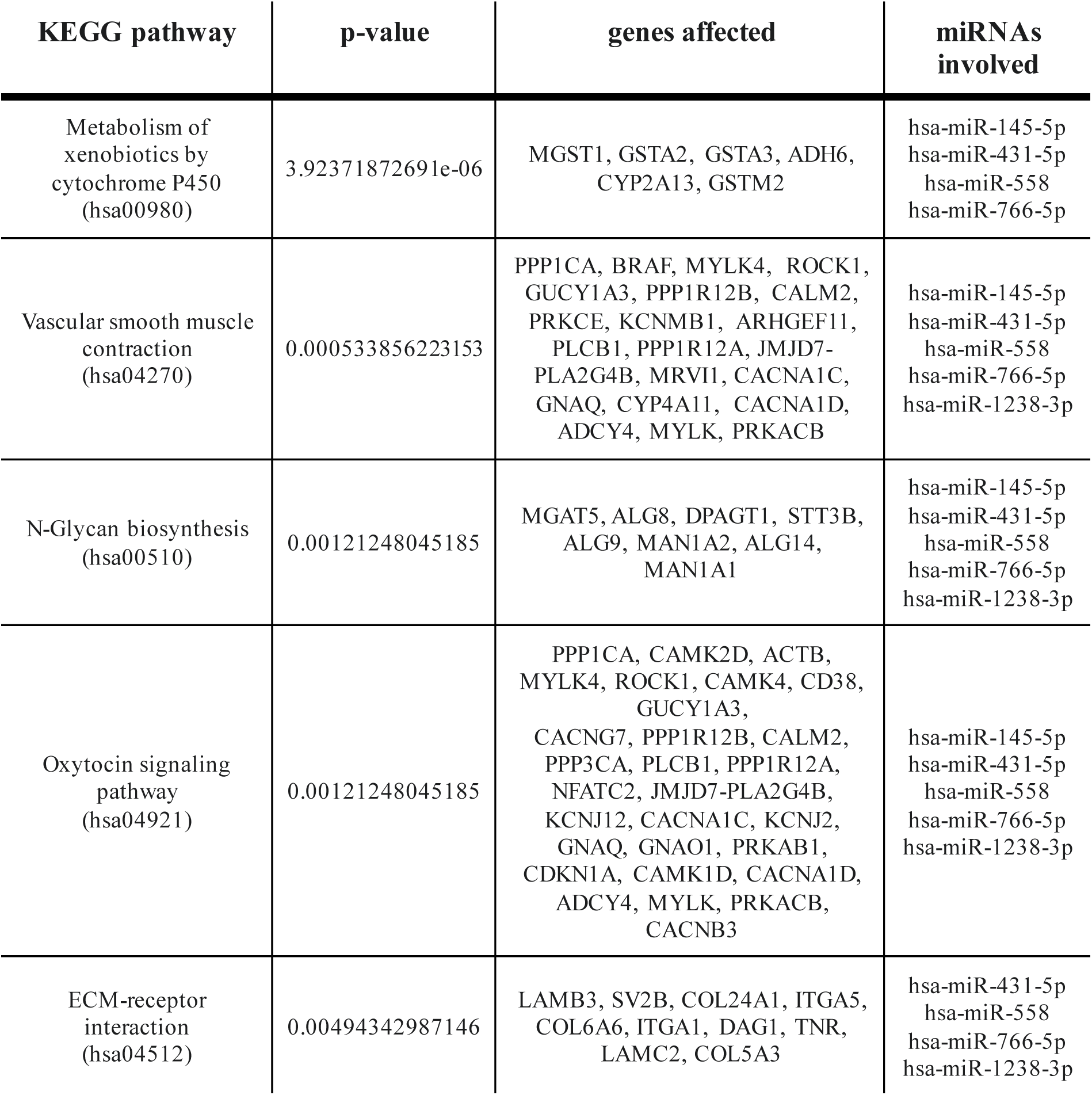
Results of DIANA-mirPath v3.0 analysis. The table shows the signaling pathways sensitive to the miRNAs we selected. Only signaling pathways for which the correlation p-value is lower than 0.05 are listed.

## References

1. Xie, Q., S. Mittal, and M.E. Berens, Targeting adaptive glioblastoma: an overview of proliferation and invasion. Neuro Oncol, 2014. 16(12): p. 1575–84.

2. Choate, K.A., et al., IDH Mutations in Glioma: Molecular, Cellular, Diagnostic, and Clinical Implications. Biology (Basel), 2024. 13(11).

3. Wang, S.S., et al., Links between cancer stem cells and epithelial-mesenchymal transition. Onco Targets Ther, 2015. 8: p. 2973–80.

4. Lei, Z.N., et al., The correlation between cancer stem cells and epithelial-mesenchymal transition: molecular mechanisms and significance in cancer theragnosis. Front Immunol, 2024. 15: p. 1417201.

5. Salami, R., et al., Circular RNAs and glioblastoma multiforme: focus on molecular mechanisms. Cell Commun Signal, 2022. 20(1): p. 13.

6. Zhong, J., et al., Circular RNA encoded MET variant promotes glioblastoma tumorigenesis. Nat Commun, 2023. 14(1): p. 4467.

7. Guo, X. and H. Piao, Research Progress of circRNAs in Glioblastoma. Front Cell Dev Biol, 2021. 9: p. 791892.

8. Zhou, Q., et al., A circular RNA derived from GLIS3 accelerates the proliferation of glioblastoma cells through competitively binding with miR-449c-5p to upregulate CAPG and GLIS3. BMC Neurosci, 2022. 23(1): p. 53.

9. Xie, F., et al., CircRNA circ_POLA2 overexpression suppresses cell apoptosis by downregulating PTEN in glioblastoma. Anticancer Drugs, 2023. 34(5): p. 652–658.

10. Kang, H., et al., Downregulated CLIP3 induces radioresistance by enhancing stemness and glycolytic flux in glioblastoma. J Exp Clin Cancer Res, 2021. 40(1): p. 282.

11. Chowdhury, T., et al., A glioneuronal tumor with CLIP2-MET fusion. NPJ Genom Med, 2020. 5: p. 24.

12. Conrad, T. and U.A. Orom, Cellular Fractionation and Isolation of Chromatin-Associated RNA. Methods Mol Biol, 2017. 1468: p. 1–9.

13. Song, X., et al., Circular RNA profile in gliomas revealed by identification tool UROBORUS. Nucleic Acids Res, 2016. 44(9): p. e87.

14. Robinson, M.D., D.J. McCarthy, and G.K. Smyth, edgeR: a Bioconductor package for differential expression analysis of digital gene expression data. Bioinformatics, 2010. 26(1): p. 139–40.

15. Li, H., et al., Sevoflurane Regulates Glioma Progression by Circ_0002755/miR-628-5p/MAGT1 Axis. Cancer Manag Res, 2020. 12: p. 5085–5098.

16. Xiao, B., et al., Circ_CLIP2 promotes glioma progression through targeting the miR-195-5p/HMGB3 axis. J Neurooncol, 2021. 154(2): p. 131–144.

17. Glazar, P., P. Papavasileiou, and N. Rajewsky, circBase: a database for circular RNAs. RNA, 2014. 20(11): p. 1666–70.

18. Schulz, J.A., et al., Characterization and comparison of human glioblastoma models. BMC Cancer, 2022. 22(1): p. 844.

19. Colopi, A., et al., Impact of age and gender on glioblastoma onset, progression, and management. Mech Ageing Dev, 2023. 211: p. 111801.

20. Rabah, N., F.E. Ait Mohand, and N. Kravchenko-Balasha, Understanding Glioblastoma Signaling, Heterogeneity, Invasiveness, and Drug Delivery Barriers. Int J Mol Sci, 2023. 24(18).

21. Eisenbarth, D. and Y.A. Wang, Glioblastoma heterogeneity at single cell resolution. Oncogene, 2023. 42(27): p. 2155–2165.

22. Mohammed, S., M. Dinesan, and T. Ajayakumar, Survival and quality of life analysis in glioblastoma multiforme with adjuvant chemoradiotherapy: a retrospective study. Rep Pract Oncol Radiother, 2022. 27(6): p. 1026–1036.

23. Shen, Y., et al., Mechanistic insights and the clinical prospects of targeted therapies for glioblastoma: a comprehensive review. Exp Hematol Oncol, 2024. 13(1): p. 40.

24. Wang, Z., et al., Cell Lineage-Based Stratification for Glioblastoma. Cancer Cell, 2020. 38(3): p. 366–379 e8.

25. Tanner, G., et al., IDHwt glioblastomas can be stratified by their transcriptional response to standard treatment, with implications for targeted therapy. Genome Biol, 2024. 25(1): p. 45.

26. Decraene, B., et al., Cellular and molecular features related to exceptional therapy response and extreme long-term survival in glioblastoma. Cancer Med, 2023. 12(10): p. 11107–11126.

27. Rybak-Wolf, A., et al., Circular RNAs in the Mammalian Brain Are Highly Abundant, Conserved, and Dynamically Expressed. Mol Cell, 2015. 58(5): p. 870–85.

28. You, X., et al., Neural circular RNAs are derived from synaptic genes and regulated by development and plasticity. Nat Neurosci, 2015. 18(4): p. 603–610.

29. Santer, L., C. Bar, and T. Thum, Circular RNAs: A Novel Class of Functional RNA Molecules with a Therapeutic Perspective. Mol Ther, 2019. 27(8): p. 1350–1363.

30. Seeler, S., et al., A Circular RNA Expressed from the FAT3 Locus Regulates Neural Development. Mol Neurobiol, 2023. 60(6): p. 3239–3260.

31. Hollensen, A.K., et al., circZNF827 nucleates a transcription inhibitory complex to balance neuronal differentiation. Elife, 2020. 9.

32. Cui, N., M. Hu, and R.A. Khalil, Biochemical and Biological Attributes of Matrix Metalloproteinases. Prog Mol Biol Transl Sci, 2017. 147: p. 1–73.

33. Hamidi, H. and J. Ivaska, Every step of the way: integrins in cancer progression and metastasis. Nat Rev Cancer, 2018. 18(9): p. 533–548.

34. Kechagia, J.Z., J. Ivaska, and P. Roca-Cusachs, Integrins as biomechanical sensors of the microenvironment. Nat Rev Mol Cell Biol, 2019. 20(8): p. 457–473.

35. Misiorek, J.O., et al., Context Matters: NOTCH Signatures and Pathway in Cancer Progression and Metastasis. Cells, 2021. 10(1).

36. Gao, Y., et al., Circular RNA profiling reveals circRNA1656 as a novel biomarker in high grade serous ovarian cancer. Biosci Trends, 2019. 13(2): p. 204–211.

37. Garcia-Gutierrez, L., M.D. Delgado, and J. Leon, MYC Oncogene Contributions to Release of Cell Cycle Brakes. Genes (Basel), 2019. 10(3).

38. Iwadate, Y., Epithelial-mesenchymal transition in glioblastoma progression. Oncol Lett, 2016. 11(3): p. 1615–1620.

39. Liu, X., et al., Overview of the molecular mechanisms of migration and invasion in glioblastoma multiforme. J Chin Med Assoc, 2021. 84(7): p. 669–677.

40. Fontana, R., A. Mestre-Farrera, and J. Yang, Update on Epithelial-Mesenchymal Plasticity in Cancer Progression. Annu Rev Pathol, 2024. 19: p. 133–156.

41. Youssef, K.K., et al., Two distinct epithelial-to-mesenchymal transition programs control invasion and inflammation in segregated tumor cell populations. Nat Cancer, 2024. 5(11): p. 1660–1680.

42. Chen, Z., et al., Hypoxic microenvironment in cancer: molecular mechanisms and therapeutic interventions. Signal Transduct Target Ther, 2023. 8(1): p. 70.

43. Joseph, J.V., et al., Hypoxia enhances migration and invasion in glioblastoma by promoting a mesenchymal shift mediated by the HIF1alpha-ZEB1 axis. Cancer Lett, 2015. 359(1): p. 107–16.

44. Berezovsky, A.D., et al., Sox2 promotes malignancy in glioblastoma by regulating plasticity and astrocytic differentiation. Neoplasia, 2014. 16(3): p. 193–206, 206 e19-25.

